# The antiviral activity of licensed therapeutics against Mpox clade Ib, *in vitro*; alternative options for the treatment of Mpox

**DOI:** 10.1101/2025.01.11.632516

**Authors:** Amanda Horton, Helen Berryman, Yasmin M. Surani, Kevin Bewley, Matthew E. Wand, J. Mark Sutton, Julia A. Tree

## Abstract

Clade Ib mpox is a newly emerged strain of the mpox virus (MPXV). The antiviral efficacy of 12 different therapeutic drugs was evaluated, *in vitro* using a live-virus, foci reduction assay, against MPXV clade Ib. We report that antiviral activity is retained against clade Ib with inhibitory concentrations (required to reduce the viral foci count by 50% (IC_50_)) of 0.025 ± 0.018 and 43.8 ± 15.2 μM for tecovirimat and cidofovir, respectively. These values are not significantly different from those observed for clade IIb, when measured in the same foci reduction assay (IC_50_ values of 0.010 ± 0.02 and 15.7 ± 14.3 μM for tecovirimat and cidofovir, respectively). Activity was also demonstrated for other antivirals, with the IC_50_ of the active metabolites of molnupiravir (EIDD-1931; 4.47 ± 1.72 μM) and remdesivir (GS-441524; 11.8 ± 6.43 μM) and with other licensed antivirals such as ribavirin (34.4 ± 10.3 μM) and baloxavir marboxil (22.6 ± 10.5 μM). In contrast, no inhibitory activity was observed with acyclovir, L-valacyclovir hydrochloride or favipiravir (IC_50_ >100 μM). Interestingly, the anti-parasitic drugs nitazoxanide, mefloquine hydrochloride and chloroquine diphosphate, showed inhibitory activity against the clade Ib virus, with IC_50_ values of 14.5 ± 3.41, 5.37 ± 1.37 and 24.7 ± 2.38 μM, respectively. This study shows that several therapeutics, including several licensed antivirals, may offer alternative treatment options for mpox clade Ib.

---

Clade Ib mpox is a newly emerged strain of the mpox virus (MPXV) and first appeared in the Democratic Republic of the Congo (DRC) in September 2023. Due to the rapid rise in the number of cases in Africa and the potential for global spread, the World Health Organization (WHO) declared it a public health emergency of international concern (PHEIC) on August 14, 2024. Imported cases of mpox clade Ib have occurred in several countries outside of Africa, including the UK. Currently there is no licensed antiviral specifically for the treatment of mpox. Tecovirimat was approved for use against smallpox by the FDA under the Animal Rule regulations in 2018, and permission for use may be granted for patients hospitalised with severe mpox disease in some countries (Lu *et al*, 2023). However, in a clinical trial in the DRC, tecovirimat did not accelerate recovery in people who were infected with clade Ib mpox according to the US National Institutes of Health (NIH) (Lenharo, 2024). Cidofovir, another antiviral licensed for use for the treatment of cytomegalovirus, has also been used to treat mpox, (Lu *et al*, 2023). The *in vitro* activity of both tecovirimat and cidofovir has been determined against various MPXV strains, and recently, in a pre-print, tecovirimat has been reported to be active clade Ib (Postal *et al*, 2024). In the present study we evaluated the *in vitro* activity of tecovirimat and cidofovir against clade Ib, alongside several other licensed drugs with potential antiviral activity against poxviridae.

A live viral foci-inhibition assay (FIA) was designed and optimised using MPXV clades Ib and IIb (supplementary file). Vero E6 cells were seeded at 2.5 x 10^5^ cells/well on the day prior to infection. On day 0, compounds were serially diluted 1:2, in duplicate, in a 96-well plate. An equal volume of media containing MPXV clades Ib and IIb were added to every well, using a multiplicity of infection (MOI) of 0.1. Immediately afterwards, 100 μl of the compound and virus mixture was then added to Vero E6 cells, previously washed with phosphate buffered saline (PBS), and then incubated at 37ºC for 1 h to allow the virus to adsorb into the cells. Afterwards the supernatant was removed and a 1% carboxymethyl cellulose (CMC) overlay containing the same concentrations of the compounds was added. After incubation at 37ºC for 22 h the plates were fixed with 8% (wt/vol) formaldehyde. The virions were immunostained and the number of foci were counted using an ImmunoSpot S6 Ultra-V analyser with a BioSpot counting module (Cellular Technologies, Europe). An internal positive control (molnupiravir) was run on each plate to ensure consistency between assays. A no-virus control (n=2) and a virus only control (VOC) (n=10) were also included on each plate. A minimum of three independent repeats were performed for each compound. The concentration of the compound that caused a 50% reduction in foci number (IC_50_) compared to the virus only control (VOC) was calculated using a non-linear fit analysis in GraphPad Prism v10.0.3 software. A cytotoxicity assay using Vero E6 cells and Cell Titre Glo 2 (Promega, UK) was also performed to determine the cytotoxic concentration (CC_50_) of each compound. The study also assessed a focused panel of licensed antivirals against clade IIb to help evaluate any differences between the two clades and to act as a bridge between this FIA and other published studies, on clade IIb, using different methodologies.

The results, in Table 1 show the IC_50_ values generated by examining the reduction in the number of viral foci, following exposure to different compounds. For tecovirimat, here we report IC_50_ values of 25 and 10 nM against clade Ib and IIb respectively. This supports the observations made by Postal *et al* 2023, where IC_50_ values of 25.8 nM for clade Ib and 15.3 nM for clade IIb, using U2OS cells, were reported in a pre-print. Others have stated IC_50_ values for different MPXV isolates against tecovirimat for example 2.8 nM for clade I (Zr-599) (Akazawa *et al*, 2023) and 2-7.5 nM for 18 clinical mpox isolates from the 2022 outbreak (Nunes *et al*, 2023). Here we also show antiviral activity with cidofovir, *in vitro*, against clade Ib and IIb with IC_50_ values 43.8 and 15.7 μM respectively. Prevost *et al* (2024) reported lower IC_50_ values, using Vero E6 cells, for cidofovir against different strains of MPXV eg 5.2 μM (Zaire 1979 clade I), 4.21 μM (USA2033 clade IIa) and 3.8 μM (SP2833 clade IIb), although an extended seven-day plaque assay was used here and this might account for the differences seen. Others report IC_50_ values of 13.7 μM against clade I (Zr-599) (Akazawa *et al*, 2023) and 41.7 μM against MPXV B.1-China-C-Tan-CQ01 (Huo *et al*, 2024); both used Vero E6 cells with different assay methodologies. Overall, the alignment of the clade Ib IC_50_’s generated here, *in vitro*, with published data for both tecovirimat and cidofovir appears encouraging. Although, this does not explain why tecovirimat is not improving clinical outcome in humans, either in terms of clinical resolution or pain, as reported in recent reports from the STOMP clinical trial (NIH, 2024). Additionally, the potential for drug resistance to develop should also not be overlooked, especially for tecovirimat (Smith *et al*, 2023).

**Table 1.** Efficacy of different therapeutics against MPXV clade Ib and IIb, in vitro.

| **Drug name** | **MPXV Clade Ib**<br>**Geometric Mean**<br>**IC<sub>50</sub> μM ± 1SD** | **MPXV Clade IIb**<br>**Geometric Mean**<br>**IC<sub>50</sub> μM ± 1SD** | **Cytotoxicity<sup>&</sup>**<br>**Geometric Mean**<br>**CC<sub>50</sub> μM ± 1SD** |
| --- | --- | --- | --- |
| Acyclovir | >100 | n.d. | >100 |
| Baloxavir marboxil | 22.6 ± 10.5 | n.d. | >100 |
| Cidofovir | 43.8 ± 15.2 | 15.7 ± 14.3 | >1000 |
| Chloroquine diphosphate | 24.7 ± 2.38 | n.d. | 161 ± 67.4 |
| EIDD-1931<sup>#</sup> | 4.47 ± 1.72 | 3.15 ± 0.90 | >100 |
| Favipiravir (T705) | >100 | n.d. | >100 |
| GS-441524<sup>\$</sup> | 11.8 ± 6.43 | 19.2 ± 2.81 | >100 |
| Mefloquine hydrochloride | 5.37 ± 1.37 | n.d. | 18.6 ± 7.25 |
| Nitazoxanide | 14.5 ± 3.41 | n.d. | 54.2 ± 10.2 |
| Ribavirin | 34.4 ± 10.3 | n.d. | >500 |
| Tecovirimat | 0.025 ± 0.018 | 0.010 ± 0.02 | >100 |
| L-Valacyclovir hydrochloride | >100 | n.d. | >100 |

n.d = not done
At least 3 independent experiments were performed, per compound.
<sup>#</sup> EIDD-1931 is the active metabolite of the antiviral prodrug molnupiravir.
<sup>\$</sup> GS-441524 is the predominant active metabolite of the antiviral prodrug remdesivir.
<sup>&</sup> Cell viability was measured following treatment with drugs using the Cell Titer-Glo Luminescent Cell Viability Assay. CC<sub>50</sub> is the effective drug concentration that reached a 50% decrease in Vero E6 cell death.

The identification of alternative licensed antivirals for the treatment of mpox is a high priority. Here we show that the active forms of certain nucleoside analogues have antiviral activity against clade Ib and clade IIb, these include EIDD-1931 (the active metabolite of molnupiravir) (IC_50_ 4.47 and 3.15 μM respectively) and GS-441524 (the active metabolite of the remdesivir) (IC_50_ 11.8 and 19.2 μM respectively). The licensed nucleoside analogue ribavirin was also active with an IC_50_ 34.4 μM against clade Ib. Previous reports have shown that molnupiravir is active against clade IIb with a reported IC_50_ of 1.35 μM (Akazawa *et al*, 2023) and ribavirin is active against MPXV Zaire V79-1-005-scab (IC_50_ 24.2 μM (5.9 μg/ml)) (Baker *et al*, 2003). We show for the first-time inhibitory activity of GS-441524, EIDD-1931 and ribavirin against clade Ib. In the present study, other nucleoside analogues, acyclovir, L-valacyclovir hydrochloride and favipiravir did not significantly reduce the number of viral foci. The lack of inhibitory activity by acyclovir has been noted by others previously (Baker *et al*, 2003).

Drug discovery researchers now use computational drug repurposing approaches such as machine learning, to identify new compounds for the treatment of mpox. Hashemi *et al*, 2024 predicted that baloxavir marboxil, an antiviral drug used to treat influenza virus in Japan, would also inhibit mpox. Here we show for the first time, with live viral data, that baloxavir marboxil has antiviral activity against clade Ib, with an IC_50_ value of 22.6 μM.

In the present study we tested three licensed anti-parasitic medications, mefloquine hydrochloride, nitazoxanide and chloroquine diphosphate; all three compounds inhibited clade Ib with IC_50_ values of 5.37, 14.5 and 24.7 μM respectively. Others have also noted the antiviral activity of mefloquine against clade IIb virus (5.19 μM) (Akazawa *et al* 2023) and nitazoxanide (2 μM) against vaccinia virus (Hickson *et al*, 2018). Here we show for the first time the antiviral activity of chloroquine diphosphate against MPXV clade Ib.

Overall, the results shown here indicate that tecovirimat and cidofovir still exhibit antiviral activity against the new strain of mpox, clade Ib and other licensed drugs also show antiviral activity, *in vitro*. Further work is required with human cell lines, organoid cultures, and/or *in vivo* models to further validate these observations. Even so, this rapid initial screen provides frontline information from which further studies can be built upon. Cytotoxicity assays were run for all the drugs using the same cell line (Vero E6) to understand the differential between the IC_50_ and CC_50_ values. As might be expected, the active forms of the licensed antivirals molnupiravir and remdesivir show the most potential as alternative drugs for the treatment of mpox, as these drugs showed a significant therapeutic window (high CC_50_ and low IC_50_) *in vitro*, are already licensed for use against other viruses, have a known toxicity profile and are widely used. It is possible that a combination of antivirals, such as tecovirimat and molnupiravir, may offer the most potent treatment option and this warrants further investigation.

## Supporting information

Supplementary file Horton et al

## Acknowledgements

We thank the National Institute for Biomedical Research (INRB), Democratic Republic of the Congo, for isolating and submitting MPXV clade Ib to the WHO BioHub System and we thank the WHO BioHub for supplying the virus (2024-WHO-LS-003) to the UKHSA. We also thank UKHSA colleagues, Dr Naomi Coombes for preparing and titrating the MPXV clade Ib bank. We thank Dr Steve Pullan for sequencing both viral banks and Mrs Marie Claude Asselin for preparing the Vero E6 cells.

## Disclosure statement

No potential conflict of interest was reported by the author(s).

## Funding

This work was funded by the UK Health Security Agency.

## Notes

### Competing Interest Statement

The authors have declared no competing interest.

## References

Akazawa D, Ohashi H, Hishiki T, et al. Potential Anti-Mpox Virus Activity of Atovaquone, Mefloquine, and Molnupiravir, and Their Potential Use as Treatments. J Infect Dis. 2023 Aug 31;228(5):591–603. doi: 10.1093/infdis/jiad058.

Baker RO, Bray M, Huggins JW. Potential antiviral therapeutics for smallpox, monkeypox and other orthopoxvirus infections. Antiviral Res. 2003 Jan;57(1-2):13–23. doi: 10.1016/s0166-3542(02)00196-1.

Hashemi M, Zabihian A, Hajsaeedi M, Hooshmand M. Antivirals for monkeypox virus: Proposing an effective machine/deep learning framework. PLoS One. 2024 Sep 12;19(9):e0299342. doi: 10.1371/journal.pone.0299342.

Hickson SE, Margineantu D, Hockenbery DM, et al. Inhibition of vaccinia virus replication by nitazoxanide. Virology. 2018 May;518:398–405. doi: 10.1016/j.virol.2018.03.023.

Huo S, Wu L, Huang B, Liu N, et al. Identification of the VP37 pocket of monkeypox virus as a promising target for pan-orthopoxvirus inhibitors through virtual screening and antiviral assays. Emerg Microbes Infect. 2024 Dec;13(1):2373309. doi: 10.1080/22221751.2024.2373309.

Lenharo M. Hopes dashed for drug aimed at monkeypox virus spreading in Africa. Nature. 2024 Aug;632(8027):965. doi: 10.1038/d41586-024-02694-x. PMID: 39160392.

Lu J, Xing H, Wang C, et al. Mpox (formerly monkeypox): pathogenesis, prevention, and treatment. Signal Transduct Target Ther. 2023 Dec 27;8(1):458. doi: 10.1038/s41392-023-01675-2.

NIH study find tecovirimat was safe but did not improve Mpox resolution or Pain. (2024) https://www.nih.gov/news-events/news-releases/nih-study-finds-tecovirimat-was-safe-did-not-improve-mpox-resolution-or-pain.

Nunes DDS, Higa LM, Oliveira RL, et al. In vitro susceptibility of eighteen clinical isolates of human monkeypox virus to tecovirimat. Mem Inst Oswaldo Cruz. 2023 Jul 10;118:e230056. doi: 10.1590/0074-02760230056.

Postal J, Guivel-Benhassine F, Porrot F, et al. Antiviral activity of Tecovirimat against Mpox virus clade 1a,1b, 2a and 2b. Bioarchive doi: 10.1101/2024.12.20.629622.

Prévost J, Sloan A, Deschambault Y, et al. Treatment efficacy of cidofovir and brincidofovir against clade II Monkeypox virus isolates. Antiviral Res. 2024 Nov;231:105995. doi: 10.1016/j.antiviral.2024.105995.

Smith TG, Gigante CM, Wynn NT, et al. Tecovirimat Resistance in Mpox Patients, United States, 2022-2023. Emerg Infect Dis. 2023 Dec;29(12):2426–2432. doi: 10.3201/eid2912.231146.

