## Supplementary file Horton et al for "The antiviral activity of licensed therapeutics against Mpox clade Ib, *in vitro*; alternative options for the treatment of Mpox"

### MATERIALS AND METHODS

**Viruses and cells:** MPXV clade Ib (#2024-WHO-LS-003) (GISAID: EPI_ISL_19424850) was obtained from the WHO BioHub System (this virus was isolated and supplied to the WHO BioHub by the National Institute for Biomedical Research (INRB), Democratic Republic of the Congo) and MPXV clade IIb (NCPV 2206091v) was obtained from the National Collection of Pathogenic Viruses (NCPV), Porton Down, UK. Both viruses were propagated in MA104 cells (ECACC 85102918), for three days, once, to produce working viral stocks. The titre of stocks was determined by plaque assay using Vero E6 cells (ECACC 85020206) as described previously (Tree *et al,* 2015). Samples of the viral stocks were sequenced by Illumina sequencing to confirm that no mutations had occurred upon passaging.

**Compounds:** All licensed drugs were obtained from Cambridge Biosciences, UK, except chloroquine diphosphate which was purchased from Merck, UK, and were either dissolved in DMSO or water to make 10mM stocks before being diluted in assay media (Table 1).

**Drug viral foci-inhibition-assay (FIA):** A live viral foci-inhibition assay (FIA) was designed and optimised using MPXV Ib and IIb. Vero E6 cells were seeded at 2.5 x $10^5$ cells/well on the day prior to infection. On day 0, compounds (Table 1) were serially diluted 1:2, in duplicate in a 96-well plate, in DMEM media (Life Technologies) containing 1% (vol/vol) fetal bovine serum (FBS) (Life Technologies), 100 U/ml penicillin, and 100 µg/ml streptomycin (Life Technologies) and 25 mM HEPES buffer (Merck, UK). An equal volume of media containing MPXV clade Ib or IIb was added to every well, at a multiplicity of infection (MOI) of 0.1. Immediately afterwards, 100 µl of the compound and virus mixture was added to Vero E6 cells, previously washed with phosphate buffered saline (PBS), and then the cells were incubated at 37ºC for 1 h to allow the virus to adsorb onto the cells. Following this incubation period, the supernatant was removed and a 1% carboxymethyl cellulose (CMC) overlay containing 100 U/ml penicillin and 100 µg/ml streptomycin (Life Technologies), 4% (v/v) FBS and 25 mM HEPES buffer in addition to the tested compounds at the same dilutions was added. After incubation at 37ºC for 22 h, in a humidified box, the plates were fixed (overnight) with 8% (wt/vol) formaldehdye. Viral foci were stained by adapting an immunostaining protocol developed for a SARS-CoV-2 micro-neutralisation assay (Bewley *et al* 2022), with the following modifications; PBS with 1% bovine serum albumin (Merck, UK)  was used as an antibody diluent and a Vaccinia antibody (rabbit polyclonal Vaccinia virus antibody (Abcam ab35219, UK) (1/2,000) was added (100 µl/well) in place of the anti-SARS-CoV-2 antibody. Plates were allowed to air dry before being scanned on an ImmunoSpot S6 Ultra-V analyser with BioSpot counting module (Cellular Technologies, Europe). An internal positive control (EIDD-1931) was run on each plate to ensure consistency between assays. Molnupiravir

(EIDD-1931) was selected to be the internal control, as this stable compound produced highly reproducible inactivation curves and similar $IC_{50}$s with small 95% confidence intervals when the drug was present in the adsorption and overlay steps or overlay only step (no adsorption) (data not shown). A no virus control (n=2) and a virus only control (VOC) (n=10) were also included on each plate. The concentration of the drug that caused a 50% reduction in foci number ($IC_{50}$), compared to virus only control, was determined using GraphPad Prism v10.0.3 software.  At least three independent experiments were performed. A drug cytotoxicity assay using Vero E6 cells and Cell Titre Glo 2 (Promega) was also performed, in parallel, to determine the $CC_{50}$.

**Cell Viability assay:** The cell viability assay was performed using a Cell Titer-Glo luminescent cell viability assay kit (Promega). Briefly, Vero E6 cells were seeded in 96-well plates. After 24 h, various concentrations of compounds were added to the medium. After 24 h, the plates were equilibrated at room temperature for 30 min, and 100 μL of Cell Titer-Glo reagent was added to the medium. The plates were subsequently shaken on a shaker for 2 min to induce cell lysis. After a final incubation for 10 min at room temperature, luminescence activity was quantified using a CLARIOstar Plus plate reader (BMG Labtech). The effective drug concentration that reached 50% decrease in cell death was defined as the $CC_{50}$ value.

**Statistical analysis:** Statistical analyses were carried out using Prism software (Graphpad Prism 10.0.3). Data are presented as geometric mean ± SD in all experiments. The $CC_{50}$ value was calculated by non-linear fit using GraphPad prism v10.0.3 software.

### References

Bewley KR, Coombes NS, Gagnon L, *et al.* Quantification of SARS-CoV-2 neutralizing antibody by wild-type plaque reduction neutralization, microneutralization and pseudotyped virus neutralization assays. Nat Protoc. 2021 Jun;16(6):3114-3140. doi: 10.1038/s41596-021-00536-y.

Tree JA, Hall G, Pearson G, *et al.* Sequence of pathogenic events in cynomolgus macaques infected with aerosolized monkeypox virus. J Virol. 2015 Apr;89(8):4335-44. doi: 10.1128/JVI.03029-14.

**Table 1: Information about the different compounds tested against MPXV clade Ib and IIb**

| **Drug name** | **Brand name** | **Type** | **CAS Number** |
|---|---|---|---|
| Acyclovir | Zovirax | Antiviral | 59277-89-3 |
| Baloxavir marboxil | Xofluza | Antiviral | 1985606-14-1 |
| Cidofovir | Vistide | Antiviral | 113852-37-2 |
| Chloroquine diphosphate | Aralen | Antiparasitic | 50-63-5 |
| EIDD-1931 (active metabolite of molnupiravir) | Lagevrio | Antiviral | 325_04_2 |
| Favipiravir (T705) | Avigan | Antiviral | 259793-96-9 |
| GS-441524 (active metabolite of remdesivir) | Veklury | Antiviral | 1191237-69-0 |
| Mefloquine hydrochloride | Lariam | Antiparasitic | 51773-92-3 |
| Nitazoxanide | Alinia | Antiparasitic & antiviral | 55981-09-4 |
| Ribavirin | Rebetol, Ribasphere, RibaPak, Copegus, Virazole & Moderiba | Antiviral | 36791-04-5 |
| Tecovirimat | Tpoxx | Antiviral | 869572-92-9 |
| L-Valacyclovir hydrochloride | Valtrex | Antiviral | 124832-27-5 |
